# Ecology and the diversification of reproductive strategies in viviparous fishes

**DOI:** 10.1101/442830

**Authors:** Michael Tobler, Zachary Culumber

**Author notes:** Corresponding author: Michael Tobler, Division of Biology, Kansas State University, 106 Ackert Hall, Manhattan, KS, USA.

## Abstract

Shifts in life history evolution can potentiate sexual selection and speciation. However, we rarely understand the causative links between correlated patterns of diversification or the tipping points that initiate changes with cascading effects. We investigated livebearing fishes with repeated transitions from pre- (lecithotrophy) to post-fertilization maternal provisioning (matrotrophy) to identify the potential ecological drivers of evolutionary transitions in life history. Phylogenetic comparative analyses across 94 species revealed that bi-directional evolution along the lecithotrophy-matrotrophy continuum is correlated with ecology, supporting adaptive hypotheses of life history diversification. Consistent with theory, matrotrophy was associated with high resource availability and low competition. Our results suggest that ecological sources of selection contribute to the diversification of female provisioning strategies in livebearing fishes, which have been associated with macroevolutionary patterns of sexual selection and speciation.

## Introduction

Evolutionary transitions in maternal provisioning strategies represent a primary axis of reproductive life history variation in viviparous organisms (Wourms 1981; Blackburn 1992). Shifts from an ancestral strategy of females providing all resources for embryonic development prior to fertilization (lecithotrophy) to post-fertilization provisioning (matrotrophy) have been associated with the evolution of complex physiological and morphological adaptations, including placental structures of apposed maternal and embryonic tissues that facilitate nutrient transfer (Wooding and Burton 2008). Matrotrophy and placentas have evolved repeatedly in viviparous animals, including multiple invertebrate (Campiglia and Walker 1995; Hart et al. 1997; Korniushin and Glaubrecht 2003; Korneva 2005; Ostrovsky et al. 2016) and vertebrate lineages (Steward and Blackburn 1988; Wourms et al. 1988; Wake and Dickie 1998; Wildman et al. 2006). Transitions along the lccithotrophy-matrotrophy continuum may have far reaching consequences, shaping the evolution of other traits and patterns of biological diversification (Zeh and Zeh 2000; Coleman et al. 2009; Pollux et al. 2014; Furness et al. 2015; Furness et al. 2019). What evolutionary forces shape the evolution of matrotrophy, however, remains unclear (Pollux et al. 2009).

There are two, non-mutually exclusive hypotheses that view matrotrophy as an adaptation to ecological sources of selection: *(1)* Matrotrophy has been hypothesized to reduce locomotor costs associated with pregnancy (Magnhagen 1991; Shaffer and Formanowicz 1996; Miles et al. 2000). Specifically, lecithotrophic females are expected to suffer from impaired locomotion throughout gestation, while matrotrophic females with initially small embryos should avoid such costs at least in early stages of pregnancy (Miller 1975; Thibault and Schultz 1978). (2) Resource availability may shape evolution along the lecithotrophy-matrotrophy continuum. Lecithotrophy is expected to be adaptive in environments with fluctuating resource availability (Thibault and Schultz 1978), whereas matrotrophy theoretically maximizes reproductive output when resource availability is high and stable (Trexler and DeAngelis 2003).

Livebearing fishes of the family Poeciliidae are an iconic model system for testing hypotheses about the evolution of reproductive strategies at micro- and macroevolutionary scales (Evans et al. 2011). Poeciliids have undergone remarkable diversification in levels of post-fertilization maternal provisioning, with independent origins of matrotrophy in different clades (Reznick et al. 2002; Pollux et al. 2009; Pollux et al. 2014). Research on poeciliid fishes has been instrumental for the advancement of our theoretical and empirical understanding of matrotrophy (Thibault and Schultz 1978; Pollux et al. 2009; Pires et al. 2011), and there is evidence supporting various predictions of both adaptive hypotheses of matrotrophy evolution. Pregnancy in poeciliids has been shown to be associated with locomotor costs (Plaut 2002; Ghalambor et al. 2004; Quicazan-Rubio et al. 2019), and there is evidence that matrotrophy increases streamlining (Fleuren et al. 2018) and may be favored in high predation environments requiring efficient escape responses (Gorini-Pacheco et al. 2017; Hagmayer et al. 2020). At the same time, post-fertilization maternal provisioning also responds to resource availability and female condition (Trexler 1997; Marsh-Matthews and Deaton 2006; Hagmayer et al. 2018), and matrotrophy is associated with significant costs when resources fluctuate (Pollux and Reznick 2011). However, most empirical tests of the potential adaptive function of matrotrophy have been conducted at microevolutionary scales and using just a small number of species, leaving questions about the generalizability of past findings.

Here, we used phylogenetic comparative analyses of matrotrophy evolution across 94 species spanning the family Poeciliidae. Species in this family are broadly distributed throughout the Americas, found in a wide variety of ecological contexts, and have a well-resolved phylogeny (Meffe and Snelson 1989; Hrbek et al. 2007; Reznick et al. 2017), facilitating comparative analyses that contrast hypotheses about the evolutionary origins of matrotrophy. To do so, we first characterized the evolutionary dynamics of matrotrophy evolution and then used phylogenetic path analysis to test for correlations between ecological variables and variation in matrotrophy.

## Methods

### Taxon sampling and phylogenetic framework

Our analyses included 94 species (Table S1), encompassing representatives of all major genera in the family Poeciliidae. These species span a geographic range from the eastern United States south to Argentina, including Caribbean islands (Figure S1). The phylogenetic framework used for analyses was established by previous studies with similar taxon sampling (Pollux et al. 2014; Culumber and Tobler 2017). In brief, sequences for six mitochondrial (*12S, COI, CytB, ND2, tRNAvalu*, and *tRNAleu*) and 11 nuclear genes (*Beta Actin, CCND1, ENC1, GLYT, MYH6, RAG1, Rhodopsin, RPS7, SH3PX3, T36*, and *XSRC*) were obtained from GenBank, aligned, and maximum likelihood phylogenetic analysis was conducted using RAxML-HPC version 8 (Stamatakis 2014) on the CIPRES computer cluster (San Diego State University, San Diego, CA, USA). The resulting best scoring tree was highly consistent with previously published phylogenetic hypotheses for the family Poeciliidae (Hrbek et al. 2007; Pollux et al. 2014; Reznick et al. 2017). Phylogenetic trees were time calibrated with three calibration points spanning the depth of the phylogeny (see Culumber and Tobler 2017 for details), including a primary fossil calibration associated with the split separating the outgroup (*Fundulus*) from all poeciliids (55-99 Ma; Santini et al. 2009) and a secondary fossil calibration separating *Heterandria formosa* from the genus *Poecilia* (9.3-19 Ma; Ho et al. 2016). In addition, the formation of Laguna de Catemaco (Mexico), a crater lake with several endemic species, was used as a constraint on the age of the endemic *Poeciliopsis catemaco* (0.5-2.0 Ma; Mateos et al. 2002). Even though bootstrap support values of the best scoring tree were generally strong, phylogenetic comparative methods described below were conducted across 250 trees drawn at random from the bootstrap replicates to account for phylogenetic uncertainty.

### Quantifying matrotrophy and sexual selection

Matrotrophy levels for all species included in the analysis were obtained from previously published studies (Pollux et al. 2014; Olivera-Tlahuel et al. 2015). The extent of post-fertilization maternal provisioning was quantified using the matrotrophy index (MI, ln-transformed for all analyses), which is the ratio of offspring mass at birth to the mass of the egg at fertilization (Reznick et al. 2002; Pollux et al. 2009). Offspring of lecithotrophic species typically lose 25-55% of the initial egg mass during development (MI < 0.75), while continuous nutrient transfer from mother to offspring during gestation in matrotrophic species leads to MI > 1 (Reznick et al. 2002; Pollux et al. 2009).

### Evolutionary dynamics of maternal provisioning strategies

To characterize the evolutionary dynamics of matrotrophy evolution, we conducted ancestral state reconstructions of MI using PHYTOOLS (Revell 2012). To evaluate the directionality of matrotrophy evolution (increases *vs*. decreases in MI) between each node and its descendants, we extracted trait reconstructions for each node of the tree and calculated ΔMI as the observed (tip) or inferred (node) matrotrophy values subtracted from the values of its most recent ancestral node.

### Identifying ecological correlates of variation in maternal provision strategies

To test the competing hypotheses for matrotrophy evolution, we assembled a set of relevant environmental predictor variables. According to the locomotor hypothesis, matrotrophy should be associated with environments that favor high locomotor performance, such as habitats with high predation pressure or fast water currents (Reznick et al. 2007; Gorini-Pacheco et al. 2017). Hence, predictor variables included metrics of hydrology (based on the topography of each species’ range) and predation (number of predatory fish species overlapping each species’ range). According to the resource availability hypothesis, matrotrophy should be associated with environments where resources are abundant and competition is low (Trexler and DeAngelis 2003). Hence, we quantified climate (including estimates of temporal variability in temperature and precipitation patterns in each species’ range), average net primary productivity (NPP), and competition (number of poeciliid species overlapping each species’ range). The potential effects of different environmental variables on variation in maternal provisioning strategies was evaluated with phylogenetic path analysis as outlined below.

#### Quantifying hydrology, climate, and net primary productivity

Assembly of hydrological and climate variables associated with each species’ range was based on georeferenced occurrence points. We obtained 73,398 locality points from multiple sources (http://fishnet2.net/, http://gbif.org/, primary literature), representing the known distributions for all 94 species included in our study. We first removed duplicate points and retained a maximum of 100 randomly sampled localities within the native range of each species, which is sufficient to capture environmental variation even in wide-ranging species (van Proosdij et al. 2015). We further verified that all data points for a given species were at least 1 km apart to match the spatial resolution of environmental data. Any locality that did not meet this criterion was either removed for species with <100 localities or replaced by another randomly drawn locality for species with >100 localities. For all locality records, we then extracted values for three hydrological variables (elevation, slope and compound topographic index; Hydro1k: https://lta.cr.usgs.gov/HYDRO1K/), 19 climatic variables (Worldclim: http://worldclim.org/), and an estimate of net primary productivity (https://lpdaac.usgs.gov/) at a spatial resolution of ~1 km^2^ (30 arcsec) in ArcMap version 10.2.2 (ESRI Inc, Redlands, CA, USA). For each species, we calculated the median value for all 23 variables. Climatic and hydrological variables were then subjected to separate phylogenetic principal component analyses (pPCA) using a correlation matrix, as implemented in the PHYTOOLS package in R (Revell 2012). In addition to the estimate of net primary productivity associated with each species’ range, we retained two pPCA axes accounting for 76% of variation in hydrology (Table S2) and three pPCA axes accounting for 81% of variation in climate (Table S3).

#### Quantifying competition and predation

Quantifying the actual biotic interactions for a large number of species distributed across the vast geographic scale included in this study is virtually impossible. Hence, we developed two simple, objectively quantifiable metrics to approximate levels of competition and predation. We assumed that competitive interactions for the focal species primarily occur with other species of the family Poeciliidae (Alberici da Barbiano et al. 2010; Torres-Dowdall et al. 2013) and that the intensity of competition is a function of the number of coexisting species. Hence, we first analyzed overlap of distributional ranges to characterize patterns of sympatry (defined as range overlap values greater than zero; Weber et al. 2016). We created geo-referenced distributional range maps for each species by generating a convex hull around each species’ known occurrence points (see above) using ArcMap. The resulting species-specific distributions were then intersected to determine the total number of competitor species exhibiting a range overlap with a focal species. Similarly, we created a metric estimating the levels of predation by determining the total number of piscivorous fish species exhibiting a range overlap with each focal species. To do so, we obtained 271,148 locality points (http://gbif.org/) of 7,170 species across 1,602 genera and 26 families in the superclass Osteichthyes that coincide with the distribution of poeciliids. Since distributional polygons of some focal species overlapped with marine habitats (particularly in poeciliid species occurring along the Gulf of Mexico and in both island and mainland localities), we first removed species primarily inhabiting marine environments as well as non-native species, retaining 5,019 native freshwater species in 853 genera (Table S4). To identify potential predators, we conducted a genus-level literature search of dietary habits using relevant monographs (Greenfield and Thomerson 1997; Bussing 1998; Boschung et al. 2004; Miller et al. 2005; van der Sleen and Albert 2017), supplemented by the primary literature when necessary. We retained 131 genera that included species with evidence for piscivory (867 species; 73,421 locality points). Values for the number of competitors and predators were square-root-transformed prior to analyses.

#### Analytical framework

We investigated hypotheses about the hierarchical relationships among abiotic and biotic environmental factors and matrotrophy using phylogenetic path analysis as implemented in the R package PHYLOPATH (van der Bijl 2018). We developed 18 models based on *a priori* hypotheses about the effects of hydrology, climate, NPP, competition, predation, and interactions between predictor variables relevant in the context of the locomotor cost and resource limitation hypotheses of matrotrophy evolution (Figure S2). As with other phylogenetic analyses described above, path analyses were run across 250 random trees. PHYLOPATH implements model selection with covariance inflation criterion, CIC_c_ (Rodriguez 2005). Models with an average ΔCIC_c_ < 2 were considered equally supported (Burnham and Anderson 2002). Joint effects of net primary productivity and competition were visualized using non-parametric thin-plate spline regression to create a surface of matrotrophy variation (Arnold 2003). Estimation of matrotrophy surfaces was performed using the FIELDS package in R, with smoothing parameter λ = 0.005 (Nychka et al. 2007).

## Results

### Evolutionary dynamics of matrotrophy evolution during diversification of poeciliid fishes

Ancestral state reconstructions (ASR) were used to compare inferred ancestral states of matrotrophy to variation in matrotrophy observed in extant taxa (Figure 1A). ASRs across 250 trees demonstrated that lecithotrophy is not the ancestral provisioning strategy (Figure 1B). The inferred ancestral state was clearly toward the matrotrophic end of the spectrum (with a net weight gain during development) and distinctly above the levels of post-fertilization provisioning observed in most extant taxa (Figure 1B). This does not mean that matrotrophy evolved prior to lecithotrophy, but rather that the common ancestor of extant poeciliids had already evolved some degree of post-fertilization provisioning. Examining the direction of shifts in post-fertilization provisioning strategies between all nodes and their descendants revealed that reductions of post-fertilization provisioning were just as common as increases in matrotrophy (Figure 1C).

**Figure 1.**
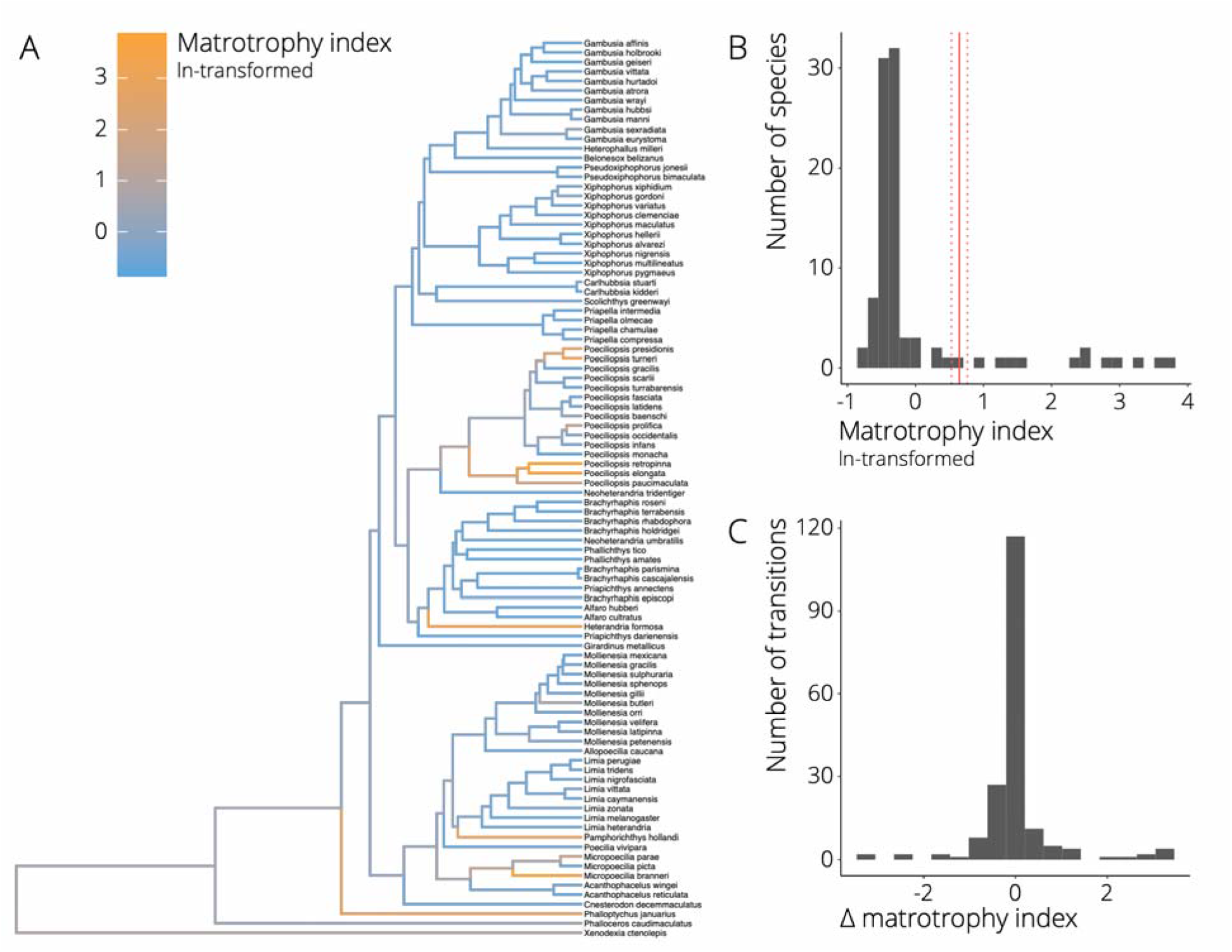
A. Best-scoring maximum likelihood tree of 94 species in the family Poeciliidae. The ancestral state reconstruction of matrotrophy is mapped onto the phylogeny, with blue colors depicting lecithotrophy and orange colors matrotrophy (as indicated by the color scale of In-transformed matrotrophy index values). B. Frequency histogram of the distribution of matrotrophy index values in extant poeciliid species. The solid red line represents the average ancestral state reconstruction for the matrotrophy index across 250 random trees with dotted lines indicating the 95% confidence interval for the estimate. C. Frequency histogram depicting the relative change in matrotrophy index between all nodes and their descendants.

### Ecological correlates of matrotrophy evolution

We contrasted a series of hypotheses about the hierarchical relationships among different abiotic and biotic environmental variables and matrotrophy using phylogenetic path analysis. Model selection identified two models with average ΔCIC_c_ < 2 across the 250 trees (Figure S3), including NPP, competition, and predation as predictor variables for variation in MI (Figure 2A). Both models indicated that NPP positively correlated with competition (*r* = 0.325, 95% CI = 0.324 – 0.326 for both supported models), which in turn was negatively correlated with matrotrophy (top model: *r* = – 0.030, 95% CI = –0.034 – –0.025; secondary model: *r* = –0.077, 95% CI = −0.081 – −0.075). In addition, there was a positive relationship between NPP and matrotrophy (top model: *r* = 0.024, 95% CI = 0.022 – 0.025; secondary model: *r* = 0.023, 95% CI = 0.021 – 0.024). Simultaneously visualizing the effects of NPP and competition on matrotrophy indicated that high levels of matrotrophy occurred when NPP was high and competition was low (Figure 2B). In the best supported model, NPP was also positively correlated with predation (*r* = 0.184, 95% CI = 0.183 – 0.185), which in turn was negatively correlated with matrotrophy (*r* = −0.082, 95% CI = −0.087 – 0.078). Notably, however, the directionality of the relationship between predation and matrotrophy was opposite to the predictions of the locomotor cost hypothesis, which posits that matrotrophy should enhance locomotor performance and be favored in high-predation environments. Although the effects sizes in the path analyses were relatively small, they were significantly different from zero, indicating that ecology has played a role in matrotrophy evolution.

**Figure 2.**
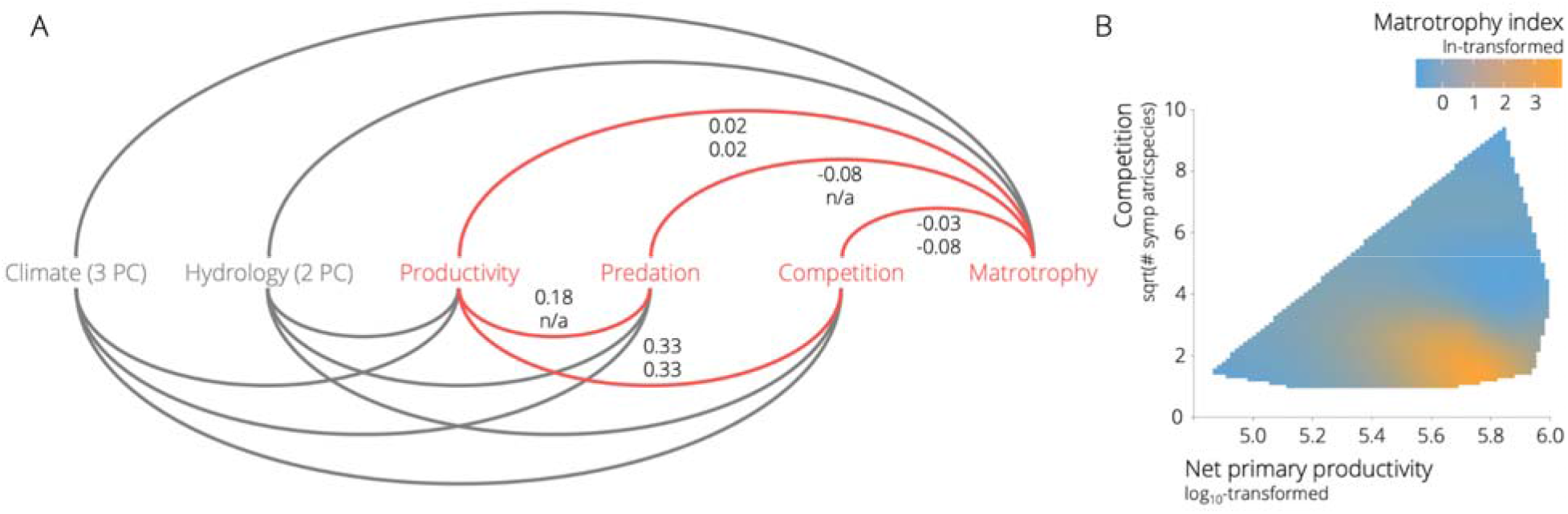
A. Representation of the full model used for phylogenetic path analysis (see Figure S2 for a comprehensive depiction of alternative models). Highlighted in red are variables and paths associated with the top models exhibiting ΔCIC_c_ < 2. Numbers represent correlation coefficients (*r*) between variables for the top model (top number) and the secondary model (bottom number). Note that the factor predation was absent from the secondary model. B. Landscape of matrotrophy variation as a function of net primary productivity and competition. Colors correspond to variation in matrotrophy, with blue colors depicting maternal provisioning strategies toward the lecithotrophy end of the spectrum and orange colors strategies toward the matrotrophy end (as indicated by the color scale of ln-transformed matrotrophy index values).

## Discussion

Shifts from pre- to post-fertilization maternal provisioning represent a major axis of life history evolution in viviparous organisms (Wourms 1981; Blackburn 1992). Even though life history evolution is generally assumed to progress from oviparity to lecithotrophic viviparity to matrotrophic viviparity (e.g., Furness et al. 2015), our analyses indicated that decreases in levels of matrotrophy were just as common as increases. Such bi-directional evolution along the lecithotrophy-matrotrophy continuum has been documented in other viviparous taxa (e.g., Dulvy and Reynolds 1997; Reznick et al. 2007) and parallels secondary losses of obligate viviparity in fishes and reptiles (Parenti et al. 2010; Recknagel et al. 2018). The standard model of linear life history evolution therefore needs reevaluation to acknowledge that the evolution of these traits is more complex and dynamic than generally appreciated.

Our analyses further suggested that ecological sources of selection are important in driving bi-directional evolution along the lecithotrophy-matrotrophy continuum. Phylogenetic path analysis identified three biotic variables that were associated with variation in matrotrophy (resource availability, competition, and predation), all of which are well-documented drivers of life history diversification in animals (Stearns 1976; Martin 1995; Wilson 2014). However, the relationship between matrotrophy and predation was opposite to the predictions of the locomotor hypothesis, even though microevolutionary analyses have indicated that matrotrophy can be favored in high predation environments requiring efficient escape responses (Gorini-Pacheco et al. 2017; Hagmayer et al. 2020). Instead, our results provided support for the resource availability hypothesis, demonstrating that high levels of matrotrophy coincided with low competition and high resource availability. This finding is consistent with the predictions of theoretical models, which predict increases in matrotrophy in environments with abundant and stable resources (Trexler and DeAngelis 2003), and empirical observations (Pollux and Reznick 2011). In addition, experimental studies have shown that maternal provisioning strategies in poeciliids respond to resource availability and female body condition (Trexler 1997; Marsh-Matthews and Deaton 2006; Hagmayer et al. 2018).

Our results largely align with previous theoretical and empirical studies; however, there are some caveats that warrant additional consideration. Most importantly, it remains to be tested whether metrics of resource availability, competition, and predation used here to facilitate continental-scale analyses accurately reflect selective regimes experienced by different species. The challenges of quantifying complex variation in biotic interactions across the vast spatial and phylogenetic scales covered in this study highlights the need for microevolutionary analyses and experimental approaches with a broader taxon sampling that allow for a better understanding of causal relationships (Culumber and Tobler 2018). For example, while there is experimental evidence for the fitness costs of matrotrophy under fluctuating resource conditions (Pollux and Reznick 2011), we still lack any empirical evidence indicating that matrotrophy provides fitness benefits over lecithotrophy under high and stable resource conditions (Pollux et al. 2009). In addition, it remains to be experimentally tested how resource availability and competition potentially interact in determining the success of different maternal provisioning strategies, especially because resource stress and competitive interactions may have non-additive effects (Hart and Marshall 2013; van Egmond et al. 2018).

## Conclusions

The role of ecological sources of selection as key drivers in life history evolution is well established at microevolutionary scales (Partridge and Harvey 1988; Reznick et al. 1990). Our study revealed that ecology also correlates with maternal provisioning strategies at macroevolutionary scales, suggesting that adaptation to resource availability and competition could explain life history diversification in livebearing fishes and potentially in other viviparous taxa (Wourms 1981; Blackburn 1992). Our findings do not exclude the possibility that other evolutionary forces are also at play. For example, variation in maternal provisioning strategies has been associated with parent-offspring conflict (Schrader and Travis 2008; Ala-Honkola et al. 2011; Pollux et al. 2014), potentially leading to antagonistic coevolution between maternal and embryonic traits that impact nutrient transfer during pregnancy (Crespi and Semeniuk 2004; Furness et al. 2015). Further studies will consequently need to explore how ecological sources of selection might interact with parent-offspring conflict to shape the evolution of matrotrophy. Disentangling the drivers of matrotrophy evolution will be particularly interesting because changes in female provisioning strategies can potentiate (or impede) sexual selection and speciation (Zeh and Zeh 2000; Zeh and Zeh 2008) and explain broad-scale patterns of diversification in viviparous taxa (Furness et al. 2019).

## Data Accessibility

Data used to conduct this study will be archived on Dryad upon acceptance of the manuscript.

## Acknowledgments

This research was supported by the National Science Foundation (IOS-1557860 to MT). We thank members of the K-State Evolution Journal Club for constructive feedback on the manuscript.

