## Supplementary appendix for "Ecology and the diversification of reproductive strategies in viviparous fishes"

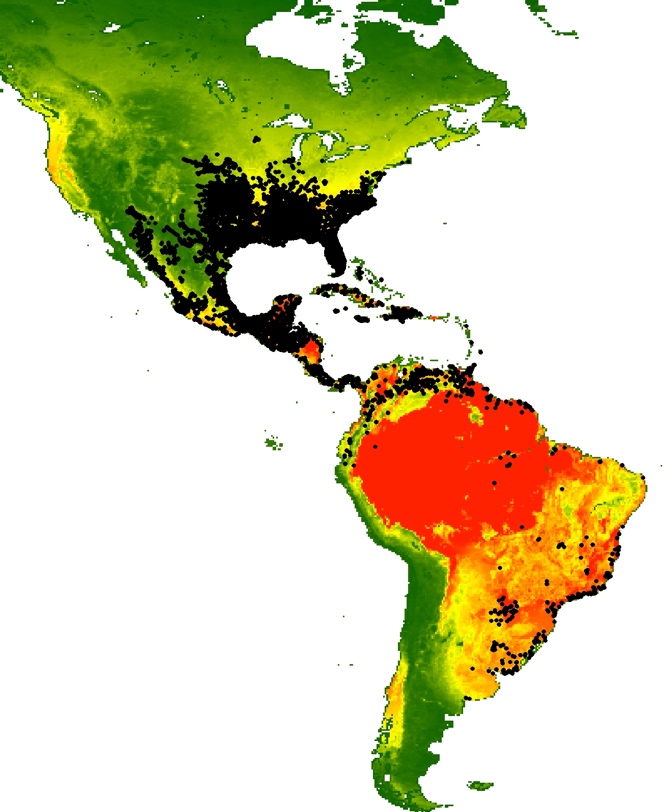

Figure S1. Map depicting the extent of the study range. Each black dot represents a collection location of one of the 94 species of poeciliids included in this study. The background coloration represents variation in net primary productivity (NPP), ranging from low values (green) to high values (red).

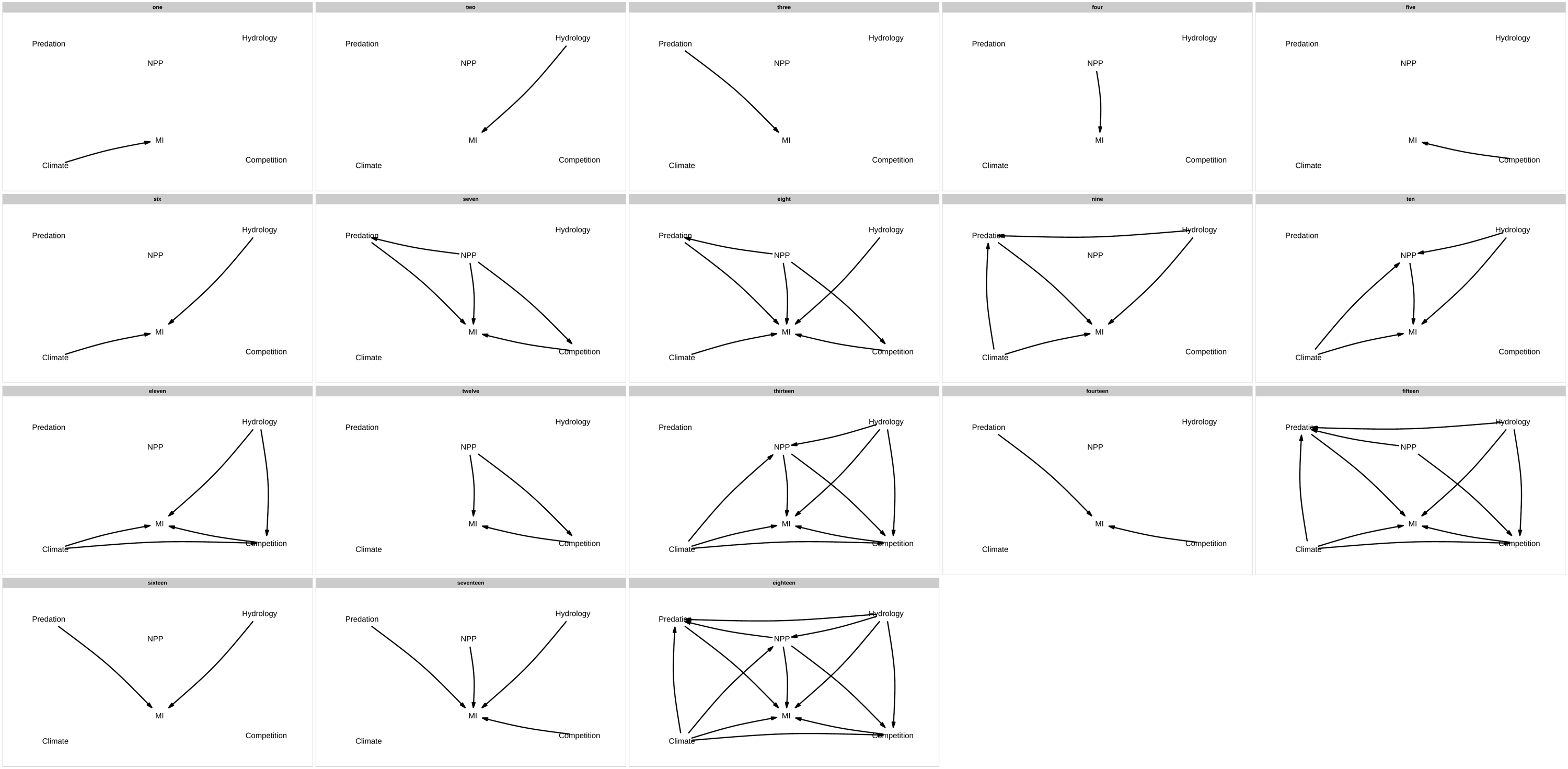

Figure S2. Alternative models describing the relationships between environmental predictors and matrotrophy: (1) Climate (3 pPC axes) affects matrotrophy; (2) hydrology (2 pPC axes) affects matrotrophy; (3) net primary productivity (NPP) affects matrotrophy; (4) competition affects matrotrophy; (5) predation affects matrotrophy; (6) abiotic environmental conditions (climate plus hydrology) affect matrotrophy; (7) biotic environmental conditions (NPP, competition, and predation) affect matrotrophy, accounting for potential effects of NPP on competition and predation; (8) abiotic and biotic conditions affect matrotrophy (combining models 6 and 7); (9) abiotic variables and predation affect matrotrophy, accounting for potential effects of abiotic variables on predation; (10) abiotic variables and NPP affect matrotrophy, accounting for potential effects of abiotic variables on NPP; (11) abiotic variables and competition affect matrotrophy, accounting for potential effects of abiotic variables on NPP; (12) NPP and competition affect matrotrophy, accounting for potential effects of NPP on competition (this specifically addresses the resource limitation hypothesis of matrotrophy evolution); (13) abiotic variables, NPP, and competition affect matrotrophy, accounting for intercorrelations of predictor variables; (14) competition and predation affect matrotrophy; (15) abiotic variables, competition, and predator affect matrotrophy, accounting for intercorrelations of predictor variables; (16) hydrology and predation affect matrotrophy (this specifically addresses the locomotor cost hypothesis of matrotrophy evolution); (17) hydrology, NPP, competition, and predation affect matrotrophy; (18) full model.

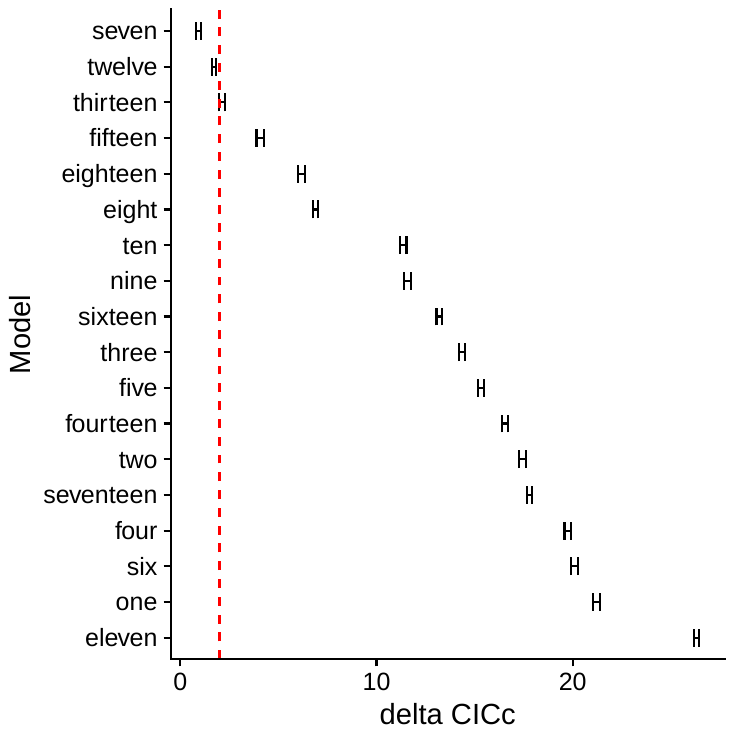

Figure S4. Contrasting different models in the phylogenetic path analysis (Fig. S2) revealed two alternative models with average ΔCICc < 2. Depicted are 95% confidence intervals of ΔCICc values for alternative models in phylogenetic path analyses conducted across 250 trees. The red dashed line represents ΔCICc = 2.

Table S1. Species included in comparative phylogenetic analyses.

*Alfaro cultratus*

*Alfaro hubberi*

*Belonesox belizanus*

*Brachyrhaphis cascajalensis*

*Brachyrhaphis episcopi*

*Brachyrhaphis holdridgei*

*Brachyrhaphis parismina*

*Brachyrhaphis rhabdophora*

*Brachyrhaphis roseni*

*Brachyrhaphis terrabensis*

*Carlhubbsia kidderi*

*Carlhubbsia stuarti*

*Cnesterodon decemmaculatus*

*Gambusia affinis*

*Gambusia atrora*

*Gambusia eurystoma*

*Gambusia geiseri*

*Gambusia holbrooki*

*Gambusia hubbsi*

*Gambusia hurtadoi*

*Gambusia manni*

*Gambusia sexradiata*

*Gambusia vittata*

*Gambusia wrayi*

*Girardinus metallicus*

*Heterandria formosa*

*Heterophallus milleri*

*Neoheterandria tridentiger*

*Neoheterandria umbratilis*

*Phallichthys amates*

*Phallichthys tico*

*Phalloceros caudimaculatus*

*Phalloptychus januarius*

*Poecilia* (*Acanthophacelus*) *reticulata*

*Poecilia* (*Acanthophacelus*) *wingei*

*Poecilia* (*Allopoecilia*) *caucana*

*Poecilia* (*Limia*) *caymanensis*

*Poecilia* (*Limia*) *heterandria*

*Poecilia* (*Limia*) *melanogaster*

*Poecilia* (*Limia*) *nigrofasciata*

*Poecilia* (*Limia*) *perugiae*

*Poecilia* (*Limia*) *tridens*

*Poecilia* (*Limia*) *vittata*

*Poecilia* (*Limia*) *zonata*

*Poecilia* (*Micropoecilia*) *branneri*

*Poecilia* (*Micropoecilia*) *parae*

*Poecilia* (*Micropoecilia*) *picta*

*Poecilia* (*Mollienesia*) *butleri*

*Poecilia* (*Mollienesia*) *gillii*

*Poecilia* (*Mollienesia*) *gracilis*

*Poecilia* (*Mollienesia*) *latipinna*

*Poecilia* (*Mollienesia*) *mexicana*

*Poecilia* (*Mollienesia*) *orri*

*Poecilia* (*Mollienesia*) *petenensis*

*Poecilia* (*Mollienesia*) *sphenops*

*Poecilia* (*Mollienesia*) *sulphuraria*

*Poecilia* (*Mollienesia*) *velifera*

*Poecilia* (*Pamphorichthys*) *hollandi*

*Poecilia* (*Poecilia*) *vivipara*

*Poeciliopsis baenschi*

*Poeciliopsis elongata*

*Poeciliopsis fasciata*

*Poeciliopsis gracilis*

*Poeciliopsis infans*

*Poeciliopsis latidens*

*Poeciliopsis monacha*

*Poeciliopsis occidentalis*

*Poeciliopsis paucimaculata*

*Poeciliopsis presidionis*

*Poeciliopsis prolifica*

*Poeciliopsis retropinna*

*Poeciliopsis scarlii*

*Poeciliopsis turneri*

*Poeciliopsis turrabarensis*

*Priapella chamulae*

*Priapella compressa*

*Priapella intermedia*

*Priapella olmecae*

*Priapichthys annectens*

*Priapichthys darienensis*

*Pseudoxiphophorus bimaculata*

*Pseudoxiphophorus jonesii*

*Scolichthys greenwayi*

*Xenodexia ctenolepis*

*Xiphophorus alvarezi*

*Xiphophorus clemenciae*

*Xiphophorus gordoni*

*Xiphophorus hellerii*

*Xiphophorus maculatus*

*Xiphophorus multilineatus*

*Xiphophorus nigrensis*

*Xiphophorus pygmaeus*

*Xiphophorus variatus*

*Xiphophorus xiphidium*

Table S2: Results of phylogenetic principal component analysis on hydrological variables from the Hydro1K dataset. We present pPC axes that were retained for downstream analyses.

| **Variable** | **pPC1** | **pPC2** |
| --- | --- | --- |
| Slope | -0.900 | 0.162 |
| Flow direction | -0.163 | -0.800 |
| Elevation | -0.537 | -0.049 |
| Compound topographic index | 0.870 | 0.007 |
| Aspect | -0.021 | 0.859 |

Table S3: Results of phylogenetic principal component analysis on climatic variables from the Worldclim dataset. We present pPC axes that were retained for downstream analyses.

| **Variable** | **pPC1** | **pPC2** | **pPC3** |
| --- | --- | --- | --- |
| BIO1 (Annual Mean Temperature) | 0.769 | 0.588 | -0.171 |
| BIO2 (Mean Diurnal Range) | -0.667 | -0.052 | -0.408 |
| BIO3 (Isothermality) | 0.767 | 0.245 | 0.255 |
| BIO4 (Temperature Seasonality) | -0.879 | -0.266 | -0.321 |
| BIO5 (Max Temperature of Warmest Month) | -0.319 | 0.328 | -0.814 |
| BIO6 (Min Temperature of Coldest Month) | 0.906 | 0.357 | 0.161 |
| BIO7 (Temperature Annual Range) | -0.883 | -0.222 | -0.369 |
| BIO8 (Mean Temperature of Wettest Quarter) | 0.122 | 0.646 | -0.593 |
| BIO9 (Mean Temperature of Driest Quarter) | 0.864 | 0.367 | -0.136 |
| BIO10 (Mean Temperature of Warmest Quarter) | 0.039 | 0.568 | -0.661 |
| BIO11 (Mean Temperature of Coldest Quarter) | 0.881 | 0.448 | 0.081 |
| BIO12 (Annual Precipitation) | 0.810 | -0.494 | -0.235 |
| BIO13 (Precipitation of Wettest Month) | 0.780 | -0.276 | -0.306 |
| BIO14 (Precipitation of Driest Month) | 0.446 | -0.712 | -0.344 |
| BIO15 (Precipitation Seasonality) | -0.210 | 0.665 | -0.164 |
| BIO16 (Precipitation of Wettest Quarter) | 0.788 | -0.307 | -0.275 |
| BIO17 (Precipitation of Driest Quarter) | 0.478 | -0.719 | -0.310 |
| BIO18 (Precipitation of Warmest Quarter) | 0.514 | -0.332 | -0.492 |
| BIO19 (Precipitation of Coldest Quarter) | 0.689 | -0.447 | -0.054 |

Table S4 (provided in a separate file: “Table S4.csv”): List of genera used to quantify predation regimes for focal species. Evidence for piscivory was found in 131 of 853 genera included.
